# A short sequence in the tail of SARS-CoV-2 envelope protein controls accessibility of its PDZ Binding Motif to the cytoplasm

**DOI:** 10.1101/2023.08.29.555304

**Authors:** Benoit Neitthoffer, Flavio Alvarez, Florence Larrous, Célia Caillet Saguy, Sandrine Etienne Manneville, Batiste Boëda

**Author notes:** These authors share first authorship (Benoit Neitthoffer, Flavio Alvarez).

## Abstract

The carboxy terminal tail of the severe acute respiratory syndrome coronavirus 2 (SARS-CoV-2) envelope protein (E) contains a PDZ-binding motif (PBM) which is crucial for coronavirus pathogenicity. During SARS-CoV-2 infection, the viral E protein is expressed within the Golgi apparatus membrane of host cells with its PBM facing the cytoplasm. In this work we study the molecular mechanisms controlling the presentation of the PBM to host PDZ (PSD-95/Dlg/ZO-1) domain-containing proteins. We show that at the level of the Golgi apparatus, the PDZ-binding motif of the E protein is not detected by E C-terminal specific antibodies neither by PDZ domain-containing protein binding partner. Four alanine substitutions upstream of the PBM in the central region of the E protein tail is sufficient to generate immunodetection by anti-E antibodies and trigger robust recruitment of the PDZ domain-containing protein into the Golgi organelle. Overall, this work suggests that the presentation of the PBM to the cytoplasm is under conformational regulation mediated by the central region of the E protein tail and that PBM presentation probably does not occur at the surface of Golgi cisternae but likely at post-Golgi stages of the viral cycle.

## Introduction

Viroporins are a class of small hydrophobic viral proteins capable of forming pores in host cell membranes. These proteins, found in most RNA viruses, are often multifunctional and involved in a range of key aspects of viral life cycle, including replication and viral genome assembly, as well as entry and release of viral particles into infected cells (1). During infection, viroporins impact cellular homeostasis and affect multiple cellular processes, such as trafficking, signaling and induction of cell death by apoptosis (2). Viroporins, due to their critical role in viral infection, have emerged as an attractive target for the development of antiviral therapies (3). SARS-CoV-2 viral genome contains three viroporin encoding genes, the envelope E, the ORF3a and the ORF8, that have been shown to form active ion channels when inserted into membranes (4–7). Out of these, the Envelope protein gene stands out for its high conservation between the SARS-CoV-1 and SARS-CoV-2 viruses, with a similarity of 94.7%. On the other hand, the conservation of Orf3a is at 73% (8), and ORF8 exhibits a lower conservation level of 30% (9). The SARS-CoV-2 E protein is a 75-amino acids (aa) small integral membrane protein that has a short N-terminal luminal domain (8aa), a central hydrophobic helical transmembrane domain (30aa), and a C-terminal tail (37aa) exposed to the cytoplasm (Figure 1A). While the N-terminal part has no known function, the transmembrane domain is able to oligomerise and form a homopentameric ion channel (10–13). The C-terminal cytoplasmic tail region encodes a Golgi targeting signal (residues 50-65 aa) (14) and is implicated as well in the interaction with host proteins. In particular, the last four carboxy-terminal amino acids of E (DLLV residues) fits the consensus type II PDZ-binding motif (PBMs) sequence (-X-ϕ-X-ϕ COOH) with ϕ signifying a hydrophobic residue allowing to interact with PDZ domain containing proteins (15, 16).

**Figure 1:**
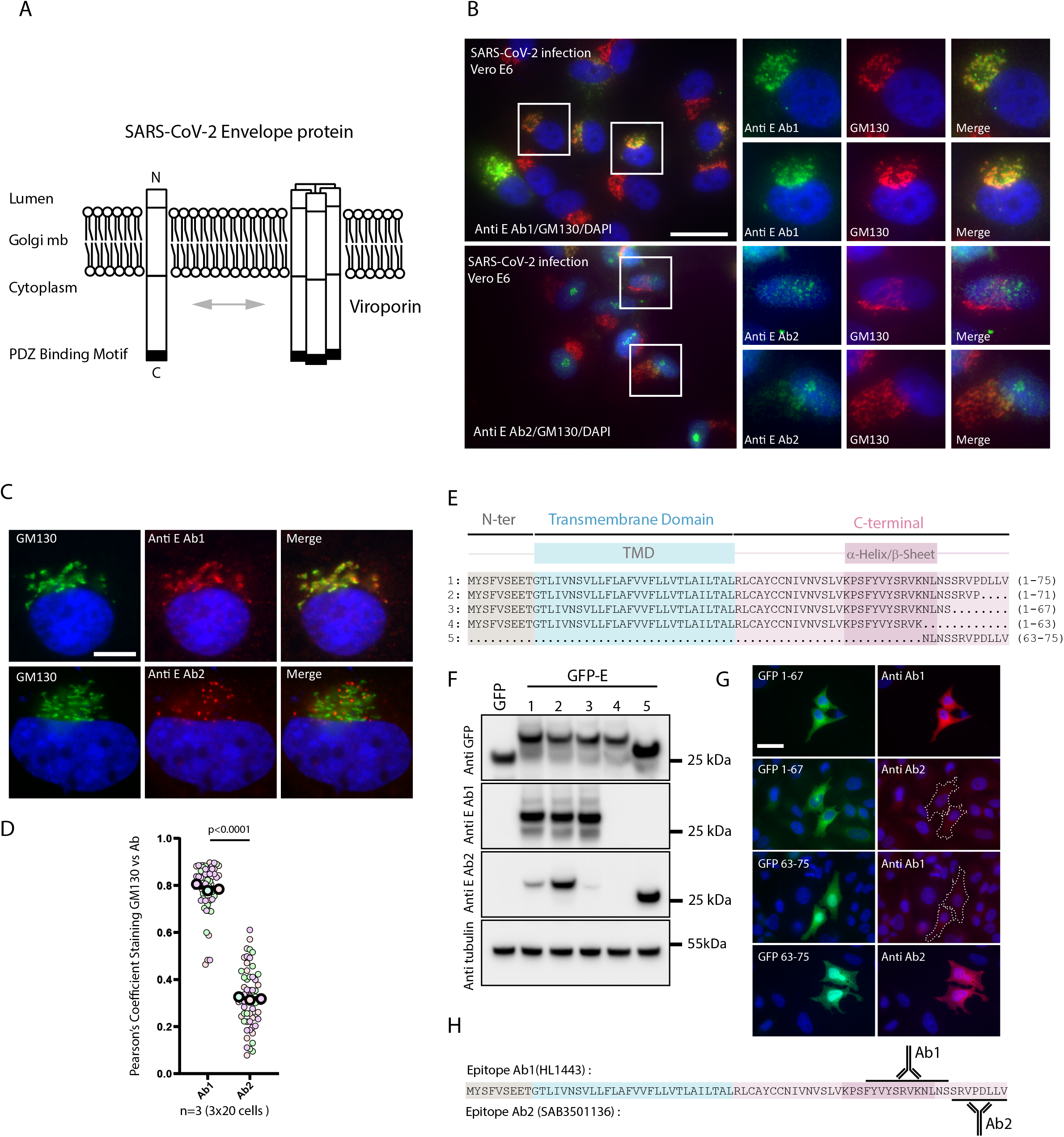
Non overlapping localizations of E protein detected by two different anti-E antibodies during infection and transfection and epitope mapping. (A) Schematic representation of E protein inserted in the host Golgi membrane. (B) Vero-E6 cells infected with SARS-CoV-2 and co-stained with Anti E Ab1 (top panel) or Ab2 (bottom panel) and GM130 (red) and DAPI (blue). Bar 30µm (C) Hela cells transfected with tagless encoding E construct and co-stained with Anti E Ab1 (top panel) or Ab2 (bottom panel) and GM130 (red) and DAPI (blue) Bar 10µm. (D) Pearson’s Correlation coefficient calculated between Anti-E (Ab1 left, Ab2 right) and GM130 staining. The p value represents the result of an unpaired two-tailed t test done on the mean of three independent experiments. (E) Graphical representation of E protein domains (top) and protein sequence alignment of the 5 different E protein fragments used in this panel (below). (F) Western Blot analysis of E protein fragments detected by anti-GFP, anti-Ab1 (Genetex, cat#HL1443) or Ab2 (SIGMA, cat#SAB3501136) commercial antibodies. Anti-tubulin is used as loading control. (G) Immunofluorescence with Ab1 or Ab2 antibodies (red) on Hela cells transfected with GFP tagged (1-67) and (63-75) E fragment. DAPI in blue. Bar 30µm. (H) Schematic representation of the epitope detected by Ab1 and Ab2 antibodies

PDZ domain-containing proteins are involved in biological processes of particular interest in viral infection such as cell junction formation, cell polarity establishment, signal transduction, metabolism and immune system signaling (17). The perturbation of the function of cellular PDZ proteins is a widely used strategy for viruses to enhance their replication, disseminate in the host, and transmit to new hosts (18). Infections with recombinant SARS-CoV-1 viruses lacking an E protein PBM are attenuated in mice, and accompanied by a decreased expression of inflammatory cytokines during infection, and a substantial increase of survival. In contrast, all mice infected with viruses containing E protein PBM die rapidly (19). SARS-CoV-1-ΔE or ΔPBM reverted to a virulent phenotype after serial passages in mice. Interestingly, the revertant viruses displayed either a new chimeric protein containing a PBM or a restored PBM sequence on the E protein (20). These findings highlight the protein E PBM sequence as a component of the virulence of the pathogen, as well as raise the question about the specific functional role of this PBM. The association of some viral PBMs with PDZ proteins results in PDZ proteins loss of function, either through proteasome-mediated degradation or through aberrant sequestration in cellular structures (17). In infected cells or when expressed from epitope-tagged cDNA, SARS-CoV-1 E localizes at the membranes of the endoplasmic reticulum-Golgi intermediate compartment (ERGIC) and Golgi apparatus (14, 21–23). Two-hybrid screenings have identified human PDZ domain-containing proteins binding to E SARS-CoV-1 PBM sequence such as Syntenin or the tight junction (TJ) protein PALS1 (19, 24). Interestingly, it has been proposed that, by recruiting and sequestrating PALS1 to the Golgi apparatus, SARS-CoV-1 E protein disrupts its trafficking to the tight junction, leading to progressive disassembly of the TJ and leakage between adjacent epithelial cells, ultimately promoting infiltration of the virus into the underlying tissue (24). A screening performed with the last 12 residues of E SARS-CoV-2 proteins has recently identified new PDZ containing proteins interactors such as the TJ protein ZO-1 as well as LNX2, PARD3, and MLLT4 proteins (16, 25). ZO-1 has three PDZ domains but only the second one binds to the PBM motif of the E protein of SARS-CoV-2 (16, 26). ZO-1 localization has been shown to be perturbed during SARS-CoV-2 infection (27) but the role of the E protein in this delocalization remains unclear. In this study, we determined SARS-CoV-2 E intracellular localization during infection and transfection and studied the molecular mechanism controlling the PBM presentation to the cytoplasm upon E protein expression in epithelial cells.

## Results

### Immunodetection of SARS-CoV-2 E protein tail during infection and transfection

To clarify the E protein localization pattern during infection, immunofluorescences were performed on Wuhan SARS-CoV-2 Vero-E6 infected cells. In these assays, two novel commercial antibodies (Ab1 [HL1443] from Genetex) and Ab2 ([SAB3501136] from SIGMA), both generated in rabbits against an unspecified C-terminal peptide, were employed. Ab1 co-localizes strongly with GM130/Golgi staining (Figure 1B, upper panels); reminiscent of what was observed with SARS-CoV-1. Surprisingly, Ab2 does not, and instead displays a more vesicle-like pattern (Figure 1B, bottom panels). In a complementary approach we transfected Hela cells with tag-free E protein-encoding plasmid and performed immunofluorescence experiments using either Ab1 or Ab2 antibodies in combination with GM130 Golgi staining. Similarly, the staining of Ab1, but not Ab2, coincides with that of GM130 (Figure 1C), with Ab2 instead decorating vesicle-like structures. Pearson’s correlation coefficient (PCC) calculation between the viral antibodies and GM130 staining depicts the strong difference observed between the two antibodies regarding their Golgi localisation (Figure 1D). Epitopes mapping of Ab1 and Ab2 antibodies reveals that Ab1 but not Ab2 recognises a GFP tagged E construct deleted of the last 8 aa (construct 3, 1-67 aa) while Ab2 but not Ab1 detects a GFP construct fused to the last 12 aa of the E protein (construct 5, 63-75 aa) in Western blotting and Immunofluorescence assays (Figure 1 E-G). This shows that Ab1 and Ab2 have distinct epitope binding sequences on the E protein and that the Ab2 recognizes the extreme end of the E protein while Ab1 a more upstream epitope (Figure 1H). Overall, this suggests that, at the level of the Golgi apparatus, the last residues of the Envelope protein encompassing the PBM are not presented to the cytoplasm in a way compatible with immunodetection.

### Lysosomal compartmentalization enhances accessibility of E-protein carboxy-terminal extremity

To explore the distinct locations of the E protein, N-terminally hemagglutinin (3xHA)-tagged SARS-CoV E protein was cloned to perform dual staining. Similar to the findings from prior experiments utilizing anti-Ab1 antibodies, anti-HA E antibodies stains the E Golgi pool. Conversely, anti-Ab2 antibodies stain peri-Golgi vesicles (Figure 2A). HA staining with Lysotracker, a cell-permeable fluorescent dye that stains acidic compartments within a cell, such as lysosomes, indicates that the majority of vesicles stained by the Ab2 antibody correspond to lysosomes (Figure 2B). Protein extract of Hela cells transfected with HA-tagged E protein-encoding plasmid were further analyzed by western blotting experiments using anti-HA, anti-Ab1 and anti-Ab2 antibodies (Figure 2C). While an expected ≃15 kDa band is detected using the anti-HA antibody, two bands migrating at 15 and 10 kDa are detected with Ab1 and only one band migrating at 10 kDa with Ab2. From these results we hypothesized that portions of the E protein pool are undergoing proteolysis cleavage leading to the clipping of the N terminal HA-tag (5 kDa) as shown by the detection of the 10 kDa band by Ab1 and Ab2 but not by anti-HA antibodies. HA-tag cleavage by endogenous proteases activated by the lysosomal compartment has already being reported previously (28, 29). More surprisingly, the 15 kDa band is not detected by the Ab2 antibody, suggesting that, in the HA-tagged full-length E protein, the accessibility of the C-terminal extremity, which includes the PBM, is compromised. However, this epitope is detected in the band at 10 kDa, suggesting that lysosomal compartmentalization improves the accessibility of the last amino acids of the E protein to Ab2 antibody. In order to check the role of lysosomal activity into Ab2 immunoreactivity we treated HA-E transfected Hela Cells with various lysosomal inhibitors (Figure 2D). While DMSO treated or MβCD (cholesterol inhibitor) control Hela cells display the expected 10 kDa band, this one is absent from the cells treated with Bafilomycin, Chloroquine or NH4Cl lysosomal inhibitors preventing autophagic proteolysis. Overall, these results indicate that the accessibility of the carboxy-terminal extremity of the E requires the recruitment of the protein to the lysosomes.

**Figure 2:**
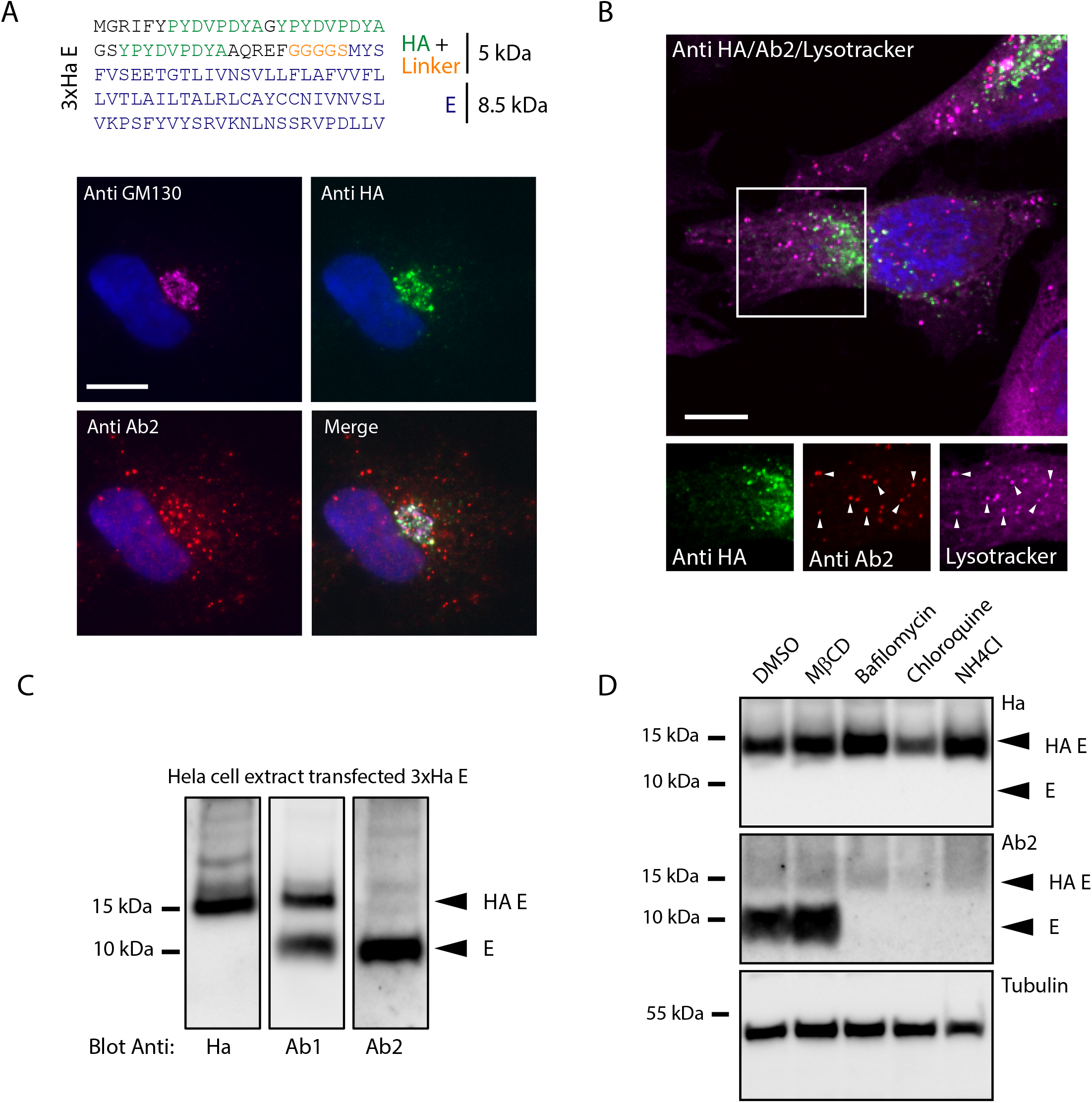
Detection of the E C-terminal epitope in the lysosome compartment. (A) Sequence of HA-E fusion construct (top). HA-E Hela cells transfected with HA-E construct and stained with anti-HA (Green), anti Ab2 (red) and anti GM130 (magenta) (bottom). (B) Hela cells transfected with HA-E construct and stained with anti-HA (green), anti-Ab2 (red) and Lysotracker (magenta). (C) Western Blot analysis of Hela cells extract transfected with HA-E construct and blot with anti-HA (right), Anti-Ab1 (middle) and Anti-Ab2 (left). (D) Western Blot analysis of Hela cells transfected with HA-E construct and treated with the indicated drugs and stained with anti-HA (top), anti-Ab2 (middle) and anti-tubulin (loading control, bottom). Bar 10µm.

### Identification of a sequence in E protein regulating the presentation of its PBM to the cytoplasm

Ab2 reduced antigenicity towards E Golgi pool could be under the influence of specific sequences within the viroporin. To explore this further, we took advantage of previous works on SARS-CoV-1 and 2 that identified and characterized several mutations within the E protein sequence (Figure 3A). These mutations affects different aspects of the viroporin biology such as its channel activity (6) (mutant 5), palmitoylation on cysteins (30) (mutant 6), potential glycosylation on aspartate residues (31) (mutant 7 and 9), a putative β-coil-β / alpha-Helix motif (11, 14) (mutant8), a putative phosphorylation (mutant 2, 3, 4 and 10) and PDZ domain binding (19) (mutant 11). HA-tagged E WT and mutant constructs were transfected in Hela cells and further analyzed by western blot using anti-HA, Ab1 and Ab2 antibodies (Figure 3B). An expected 15 kDa migrating band is observed with the anti-HA antibody in all the conditions. A 15 kDa/10 kDa doublet was detected with Ab1 antibody with the exception of mutants 8 and 9. Finally, a 10 kDa migrating band was observed using the Ab2 antibody with the exception of mutants 7 and 11. Remarkably, the FYVY to AAAA mutation (mutant 8) leads to the immunodetection of a 15 kDa band with Ab2 antibody in addition to the 10 kDa band (Figure 3B). To further demonstrate that the FYVY to AAAA mutation improves the detectability of E protein by the Ab2 antibody, we analyzed Hela cells transfected with HA-E (Figure 3C) or tag-free E (Figure 3D) encoding constructs by immunofluorescence. In these conditions, and contrarily to WT sequence, the FYVY to AAAA mutant is detected at the Golgi apparatus by the Ab2 antibody as shown by the co-staining with GM13O. Altogether, these results indicate that the FYVY to AAAA mutation enhances the accessibility of the E protein C-terminal residues to the Ab2 binding at the level of the Golgi apparatus.

**Figure 3:**
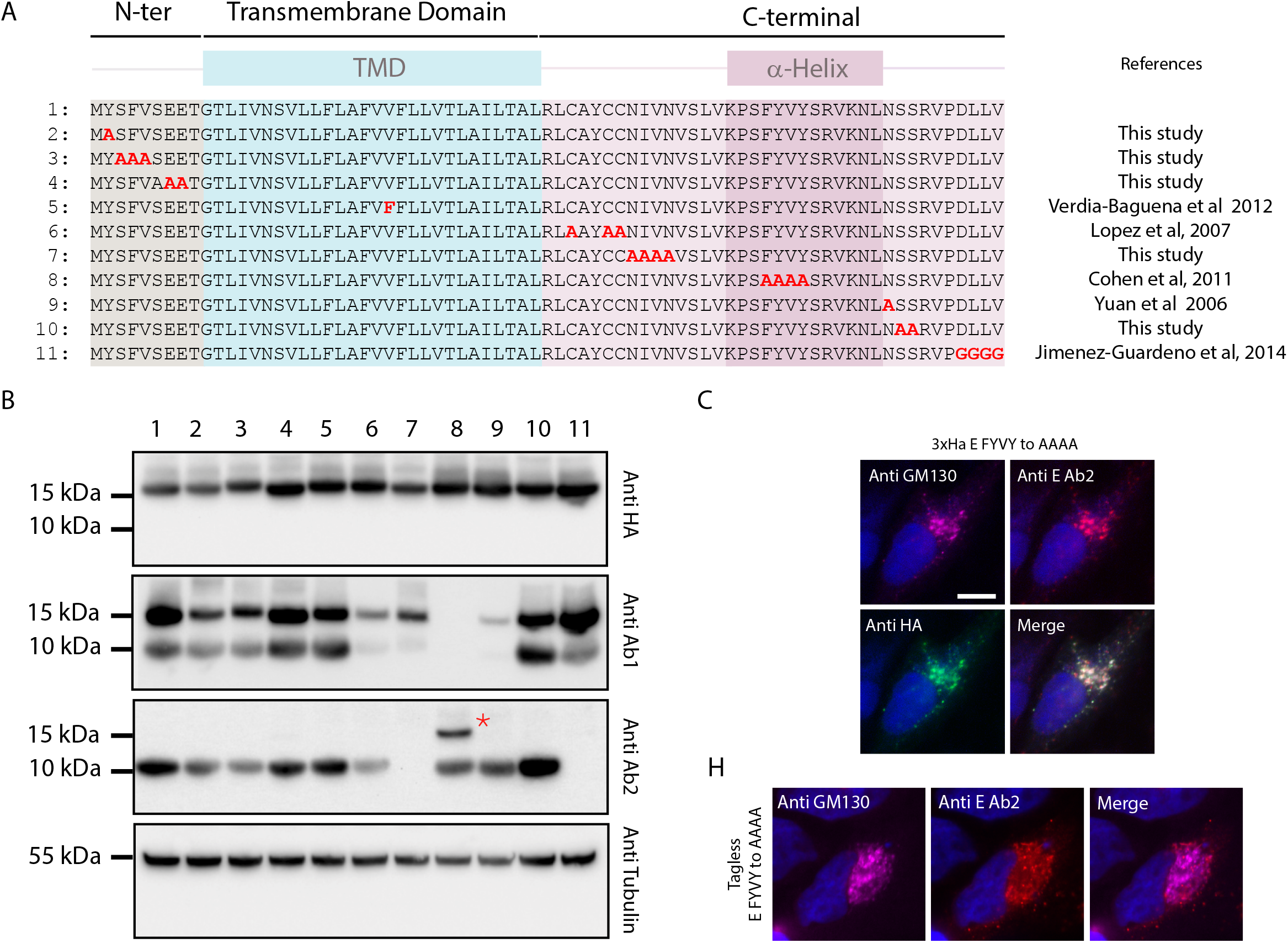
Identification of a sequence in E governing C-terminal epitope presentation. (A) Graphical representation of E protein domains (top) and protein sequence alignment of the mutant constructs used in this panel. (B) Western Blot analysis of Hela cells extract transfected with the different HA-tagged E mutant constructs and blot with anti-HA, anti-E AB1, anti-E Ab2, and anti-tubulin. The red star indicates the 15 kDa band detected with Ab2 and mutant 8. (C) Hela cells transfected with HA-E mutant 8 construct and stained with anti-GM130 (Magenta), anti-Ab2 (red), anti-HA (green) and DAPI. (D) Hela cells transfected with tagless E mutant 8 and stained with anti GM130 (magenta), Anti Ab2 (red) and DAPI. Bar 10µm.

### The FYVY sequence of the E protein governs the recruitment of ZO-1 to the Golgi apparatus

The E protein expressed at the Golgi apparatus is thought to recruit PDZ domain-containing proteins to this organelle. To test this hypothesis, Hela cells were co-transfected with GFP-ZO-1 full length construct (Figure 4A) and an Alfa tagged E WT(25, 32) encoding plasmid and Golgi localization was quantified using GM130. GFP-ZO-1 full construct exhibits limited co-localization with GM130 (Figure 4B). Because we previously observed that accessibility to the C-terminal extremity of the E protein was improved by the FYVY sequence mutation, we expressed GFP-ZO-1 together with FYVY to AAAA E mutants in Hela cells (Figure 4C). In contrast to the WT sequence, the E-FYVY to AAAA mutant robustly recruited ZO-1 protein to the Golgi apparatus (Figure 4C and D). This Golgi recruitment triggered by the E-FYVY mutant is specific to ZO-1 as it was not observed with construct expressing the full length homologue PDZ domain-containing protein ZO-2 fused to GFP (Figure 4E). Taken together, these results indicate that the FYVY sequence increases the exposure of the PDZ binding motif to the cytoplasmic compartment thus regulating the binding of the E PBM to the PDZ domain of host proteins.

**Figure 4:**
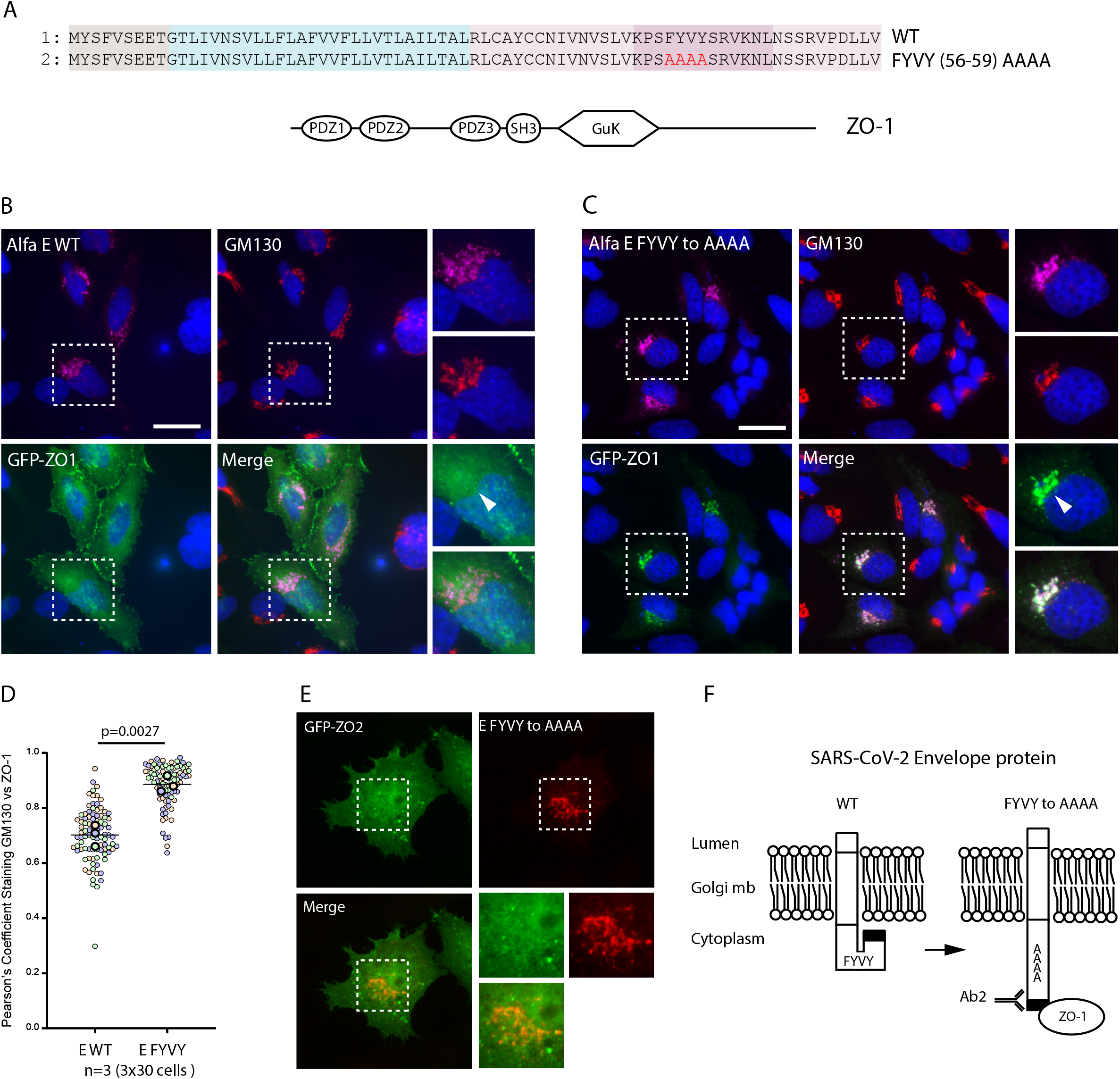
ZO-1 recruitment to E Golgi pool is regulated by the FYVY motif. (A) Sequence alignment of the WT and mutant FYVY to AAAA constructs and graphical representation of the full-length ZO-1 protein. (B) Immunofluorescence of Hela cells transfected with WT Alfa-E and full-length GFP tagged ZO-1 construct and stained with anti-Alfa and anti-GM130 antibodies. (C) Immunofluorescence of Hela Cells transfected with Alfa-E FYVY to AAAA mutant and full length GFP tagged ZO-1 construct and stained with anti-Alfa and anti-GM130 antibodies. (D) Pearson’s Correlation coefficient calculated between GFP signal and GM130 staining. Zoomed areas displayed are taken from the dashed square. The p value represents the result of an unpaired two-tailed t test done on the mean of three independent experiments. (E) Immunofluorescence of Hela Cells transfected with Alfa-E FYVY to AAAA mutant and full-length GFP tagged ZO-2 construct and stained with anti-Alfa and anti-GM130 antibodies. Bar 20µm. (F) Schematic representation of the conformational shift hypothesis induced by the FYVY to AAAA mutation and its effect on Ab2 and ZO-1 binding.

## Discussion

In this study, we investigated the intracellular localization of the E protein and its coupling to the recruitment of the PDZ containing protein ZO-1. We show that during infection and tagless transfection, the E protein is detected at the Golgi apparatus using an antibody recognizing an internal epitope (Ab1) and in the lysosomal compartment with an antibody recognizing the final C-terminal residues (Ab2). Furthermore, we showed that E protein cannot recruit the PDZ domain-containing protein ZO-1 to the Golgi apparatus. However, we established that immunodetection by Ab2 at the Golgi, as well as the Golgi sequestration of ZO-1, can be induced by introducing alanine substitutions to the FYVY motif in the E protein.

Based on our observations, one could speculate that the accessibility of the PBM sequence might potentially be controlled by either amino acid side chain modifications, selective cleavage of the C-terminal sequence, or internal conformational switch. Previous studies have shown that the interactions between PDZ domains and their PBM containing partners might be negatively modulated by phosphorylation events on serine or threonine residues located close to or within the PBM (33–35). Although we cannot totally exclude this hypothesis, we have shown that substitution of both serine 67-68 with alanine residues (mutant 10, Figure 3) did not improve E protein recognition by Ab2 antibody. Alternatively, the SARS-CoV-1 E protein has been previously shown to be palmitoylated on its cysteines 40, 43 and 44 (30) and glycosylated on the asparagine residue N66 (31). Again, mutation of the concerned residues (mutants 6 and 9, Figure 3) did not modify Ab2 immunodetection.

The second hypothesis is that a protease selectively cleaves the C-terminal tail of the E protein at the Golgi apparatus, thereby inhibiting Ab2 immunodetection or binding to PDZ domains. Furthermore, this would imply that the FYVY mutation is inhibiting the protease action, therefore allowing Ab2 immunodetection and PDZ domain binding. While this type of PBM-PDZ binding regulation by protease has never been described before, it is interesting to note that the PDZ domain containing protease Htra1 has been shown to interact *in vitro* with the E protein C-terminal PBM (16). However, no commercial Htra1 inhibitors are available to date to test this hypothesis.

Regarding the third hypothesis about a conformational switch, several observations point to conformational changes of the E protein. Indeed, some E mutants do not show the expected electrophoretic profile by SDS-PAGE in Figure 3. Some mutations (8 and 9 for Ab1 and 11 for Ab2) induce a lack of band recognition that could be rationalized by the fact that these mutations fall within the epitope binding sequence of Ab1 and Ab2 antibodies respectively. However it is not clear why the mutations 7 lead to the lack of detection of the 10 kDa band and mutation 8 to the recognition of an additional 15 kDa band, as both mutations lie outside the Ab2 binding epitope sequence. These mutations might have a strong impact on the E protein folding, hiding or exposing the PBM epitope to the Ab2 antibody even under SDS-PAGE denaturing conditions where membrane protein might be resistant to unfolding and migrate anomalously due to detergent binding (36). In addition, the FYVY to AAAA substitution in the E protein sequence (mutant 8) induced a drastic change in the recognition of E Golgi pool by Ab2 antibody or ZO-1 protein, suggesting that this region plays a conformational positive role in the presentation of the PBM to the cytoplasm (Figure 3B-D, 4C and F). Previous work done by Jaume Torres group on SARS-CoV-1 E protein has established that the C-terminal part of the E protein was a region of high structural plasticity (11). The region encompassing the 40-65 residues containing the FYVY motif has been shown to adopt either a beta-sheet or an alpha-helix structure depending on the experimental conditions used (37). Recent work suggests that the conformational switch between the beta-sheet and the alpha-helix may be regulated by local membrane curvature, with beta-sheet conformation being triggered by a low curvature membrane typically found in ERGIC or Golgi organelles and alpha-helix conformation being favored by a high-curvature membrane structure such as the ones found in vesicles, budding virus or virion envelopes (38). Interestingly, alanine substitutions of the FYVY motif are thought to increase its alpha helix propensity (37), suggesting that the FYVY to AAAA mutant present a C-terminal tail constitutively locked in an alpha helix conformation. All this previous work was carried out on artificial bilayer membranes using a truncated E protein fragment (residues 8-65) lacking the PBM, which precluded the study of variation in PBM exposure as a function of conformational changes in protein E. Our work examines the behavior of the full-length protein E *in cellulo*. Overall, our study suggests that the E protein has a higher binding affinity to PDZ domains when its C-terminal region adopts an alpha helical conformation (Figure 4E). This conformation does not appear to be met at the Golgi apparatus level, which challenges the hypothesis of Golgi sequestration, at least for the ZO-1 protein. As beta-Coronaviruses use lysosomes for egress, one could hypothesized that the PBM become exposed to the cytoplasm at later stage of the virus life cycle (39).

## Experimental Procedures

### Antibodies

The following antibodies were used in this study: Mouse anti-ZO-1 (BD Trans Lab cat#610967), anti-E antibody Ab1 (Genetex, cat#HL1443) and Ab2 (SIGMA, cat#SAB3501136), anti-GM130 (BD Trans Lab cat#610823), anti-HA (SIGMA, 11867423001), Anti-Alfa (NanoTag Biotechnologies, cat#N1502), Anti-tubulin (Abcam, cat#ab6061), Anti-MyoX (NovusBio, cat# NBP1-87748), phalloidin (Thermo Fischer Scientific, cat#A22287), Hoechst/Dapi (Thermofisher).

### Cloning

Envelope cDNA (Wuhan-hu-1 strain) was synthetized by Eurofins and cloned EcoRI/XhoI into pcDNA vector fused N-terminally to GFP, 3xHA or Alfa (MSRLEEELRRRLTE peptide) tags respectively. Envelope gene mutations were introduced into WT cDNA sequence using the Q5 Site-Directed Mutagenesis Kit (New England Biolabs). All cloned were sequenced (Eurofins).

### Viral infection

Confluent monolayer of Vero-E6 cells (ATCC #CRL-1586) was seeded one day prior to infection in Labtek. Cells were washed once with PBS and inoculated during 1h with Wuhan SARS-CoV-2 virus at MOI of 0.01, diluted in DMEM without SFV. Inoculum was then removed and DMEM without SFV medium was added in each well. After 24h of infection, cells were washed with PBS and fixed during 30 min with PFA 4%.

### Transfection and Immunofluorescence

HeLa cells were transfected using the Genejuice (Novagen) or Lipofectamine 2000 (Thermofischer) transfection reagents according to the protocol of the manufacturers. At 24 h after transfection, cells were fixed with PBS PFA 4% (SIGMA, cat#F8775) for 10 min and permeabilized in PBS Triton 0.1% for 5 min before being processed for immunofluorescence. For extracellular staining protocol, coverslips with Hela cells were incubated in PBS, on ice, with red anti-Alfa FluoTag for 30 min before being washed 3 times with PBS and fixed with 2% methanol-free PFA (Electron Microscopy Sciences, cat#157-8). Coverslips were directly mounted using Prolong Diamond Antifade mounting medium (ThermoFischer) and images acquired with a Leica DM6B microscope with a 63X objective.

### Drug treatment

Hela cells were incubated overnight with Bafilomycin A1 (300 nM, SIGMA, cat# SML1661), Chloroquine (50 µM, SIGMA, cat# C6628) and NH4Cl (10mM SIGMA, cat# A9434). Nocodazole (100nM, SIGMA, cat# M-1404) and MβCD (5µM, SIGMA, cat# C4555) and Lysotracker (100 nM, cat# L12492) were incubated 1h with the cells.

### Image analysis and statistical test

Image analysis was performed using the image processing package Fiji(40). Pearson’s correlation coefficient was calculated using the JACop pluggin 36 on 40x40µm images centered around the GM130 Staining (41). The p-value represents the result of an unpaired two-tailed t test done on the mean of three independent experiments using GraphPad Prism Software.

### Cell culture

Hela cells were maintained in DMEM supplemented with 10% FBS (Invitrogen) and penicillin (100 U/ml)-streptomycin (100 μg/ml; Invitrogen) at 37°C in 5% CO2. Vero E6 cells were maintained in DMEM 5% FBS at 37°C in 5% CO2.

## Funding

This work was supported by the « URGENCE COVID-19 » fundraising campaign of Institut Pasteur.

### Abreviation

SARS-CoV: (Severe Acute Respiratory Syndrome Coronavirus);
(PDZ (PSD-95/Dlg/ZO-1): PBM (PDZ-binding motif)

## Data Availability Statement

The data that support the findings of this study are available on request from the corresponding author, Batiste Boëda, upon reasonable request.

## Declaration of interests

The authors declare that they have no known competing financial interests or personal relationships that could have appeared to influence the work reported in this paper.

## Acknowledgements

The authors thank the Photonic Bioimaging platform of Institut Pasteur and the member of PCMC laboratory for experimental discussion.

## Notes

### Competing Interest Statement

The authors have declared no competing interest.

